## supporting figures for "Inhibition of SARS-CoV-2 viral entry *in vitro* upon blocking N- and O-glycan elaboration"

**Contents**:
Five supplemental Figures

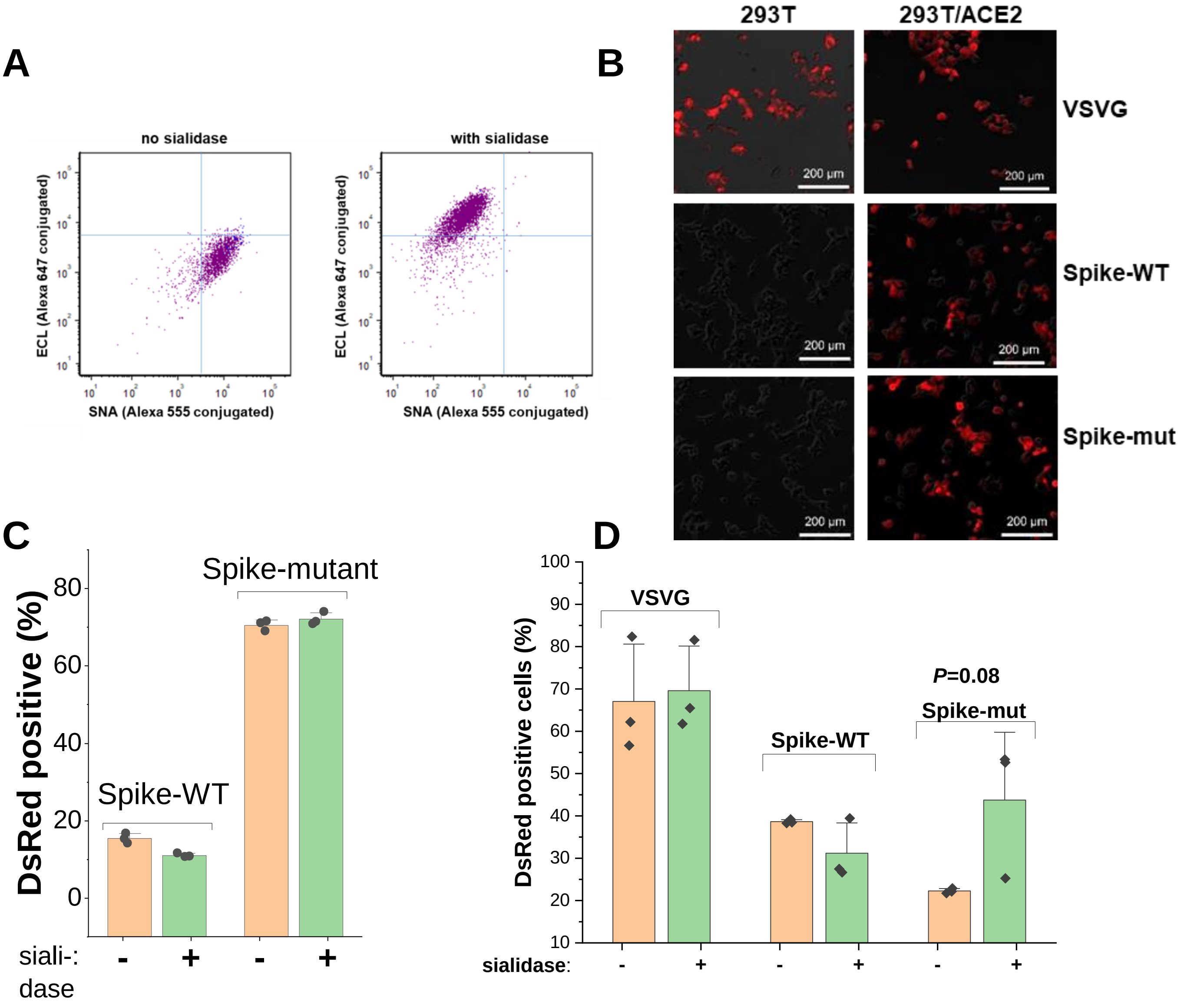


**Figure 2-figure supplement 1. Sialidase treatment studies. A.** Sialidase protocol validation. All lectins were directly conjugated with Alexa dyes. They were incubated with cells at 1-5µg/mL for 15 min before a quick wash and cytometry measurement. Compared to untreated control (left), sialidase treatment (right) decreased SNA lectin binding to α2,6 sialylated structures by 15-fold and increased ECL binding to desialylated lactosamine chains (Galβ1,4GlcNAcβ) by an order of magnitude. **B.** Pseudovirus assay. DsRed fluorescence in HEK293T and stable 293T/ACE2 cells upon addition of VSVG, Spike-WT and Spike-mutant pseudotyped virus. **C.** Sialidase treatment of pseudovirus. % DsRed positive cell data are shown for study in Fig. 2F (main manuscript). Viral entry was sialidase independent. **D.** Sialidase treatment of HEK/ACE2 cells. Pseudovirus expressing VSVG, Spike-WT and Spike-mutant were added to cells under conditions described in Fig. 2G (main manuscript). All error bars are standard deviations. Data are representative of 3 independent runs.

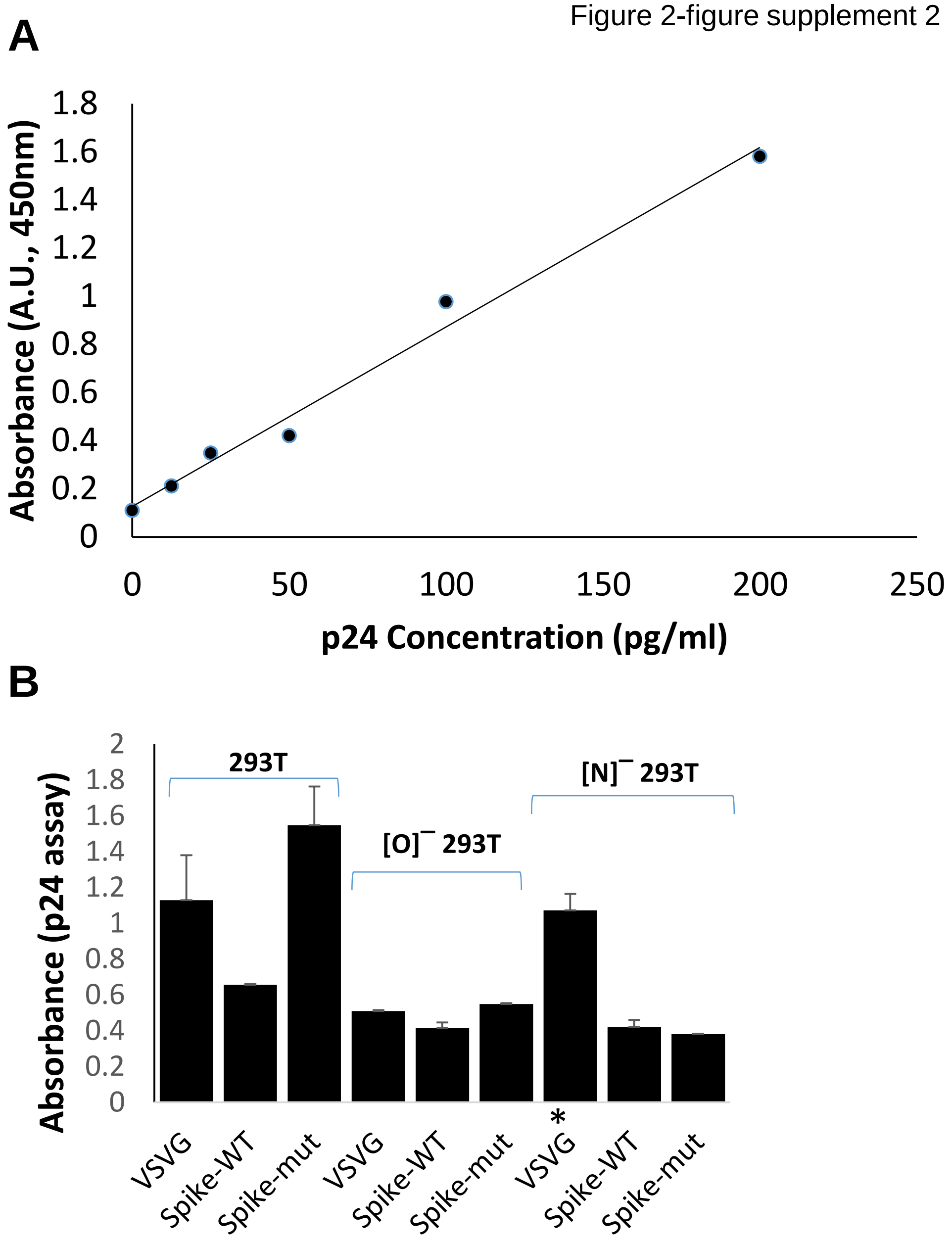


**Figure 2-figure supplement 2. p24 assay to determine viral titer. A.** Representative calibration curve for p24 standard. **B.** Absorbance value for viral stock in representative ELISA run. 9 different viruses were prepared and used in Fig. 4 (main manuscript). All virus were diluted 1:500,000 fold before ELISA measurement, except for VSVG virus made in [N]¯ 293T cells (indicated by asterisk) which were diluted 1:50,000 fold, in this instance, due to lower titer.

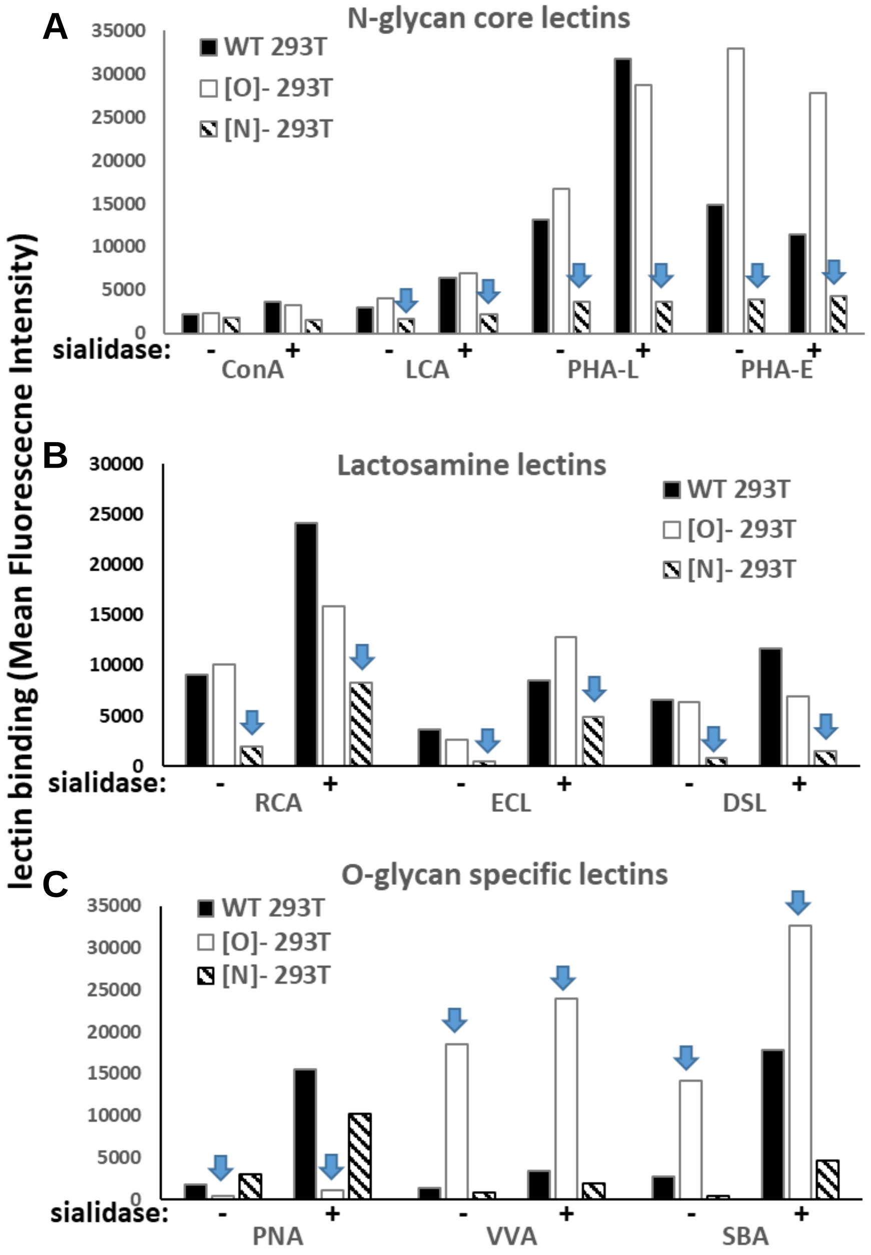


**Figure 3-figure supplement 1. Lectin binding to wild-type and glycogene-KO 293T cells.** A panel of lectins (from Vector Labs) was conjugated with Alexa dyes, either Alexa 405, 488 or 647. The binding of these fluorescent reagents to wild-type 293T, [N]¯293T and [O]¯293T cells was measured using flow cytometry. The lectins bound: **A.** N-glycan high-mannose and complex structures [ConA & LCA bind αMan in high mannose glycans; PHA-L & PHA-E bind complex glycans], **B.** lactosamine chains primarily on N-linked glycans [RCA, ECL bind terminal Gal or lactose; DSL bind β1,4GlcNAc], and **C**. O-glycan related structures [PNA binds Galβ1,GalNAc; VVA & SBA bind GalNAcα]. Measurements were made with either untreated or sialidase treated 293T cells. As seen: i. Knocking out *MGAT1* in [N]¯293T reduces lectin binding in panels A and B (see arrow). ii. Knocking out *C1GalT1* results in a dramatic decrease in PNA binding and increase in VVA and SBA binding, These data are consistent with the expected changes in lectin profile upon knocking out these N- and O-glycan specific enzymes.

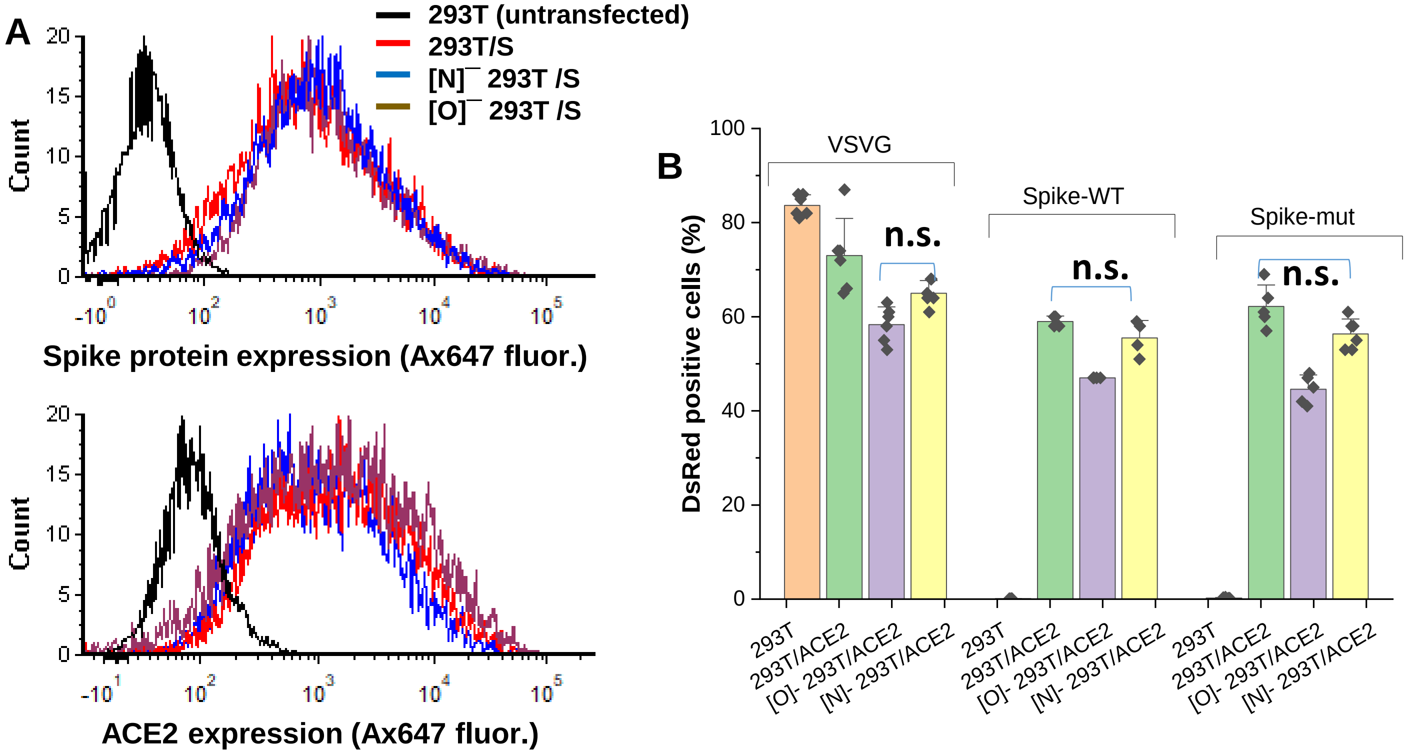


**Figure 3-figure supplement 2. Effect of ACE2 glycosylation on viral entry. A.** Surface expression of Spike-protein and ACE2. Full length Spike (top) and human ACE2 (bottom) were expressed in HEK 293T, [N]¯293T and [O]¯293T. Protein expression was measured in EGFP+ cells in the case of Spike (using anti-RBD), and on BFP+ cells in the case of ACE2 (using anti-ACE2), as these fluorescent reporters are co-expressed with surface proteins. Protein expression was comparable in all cells. Untransfected 293Ts serve as negative control. **B.** Viral entry assay. Pseudovirus expressing VSVG envelope protein, Spike-WT or Spike-mutant were added to HEK 293T cells transiently transfected to overexpress ACE2 (both wild-type 293T and glycosylation mutants). An additional control included 293T cells not expressing ACE2, which only allowed entry of VSVG pseudotyped viral particles, but not Spike bearing virus. % cells that were DsRed (reporter) positive is shown at 72h. All treatments were statistically different except as indicated by n.s. (‘not significant’). All data are from of N≥3 repeats.

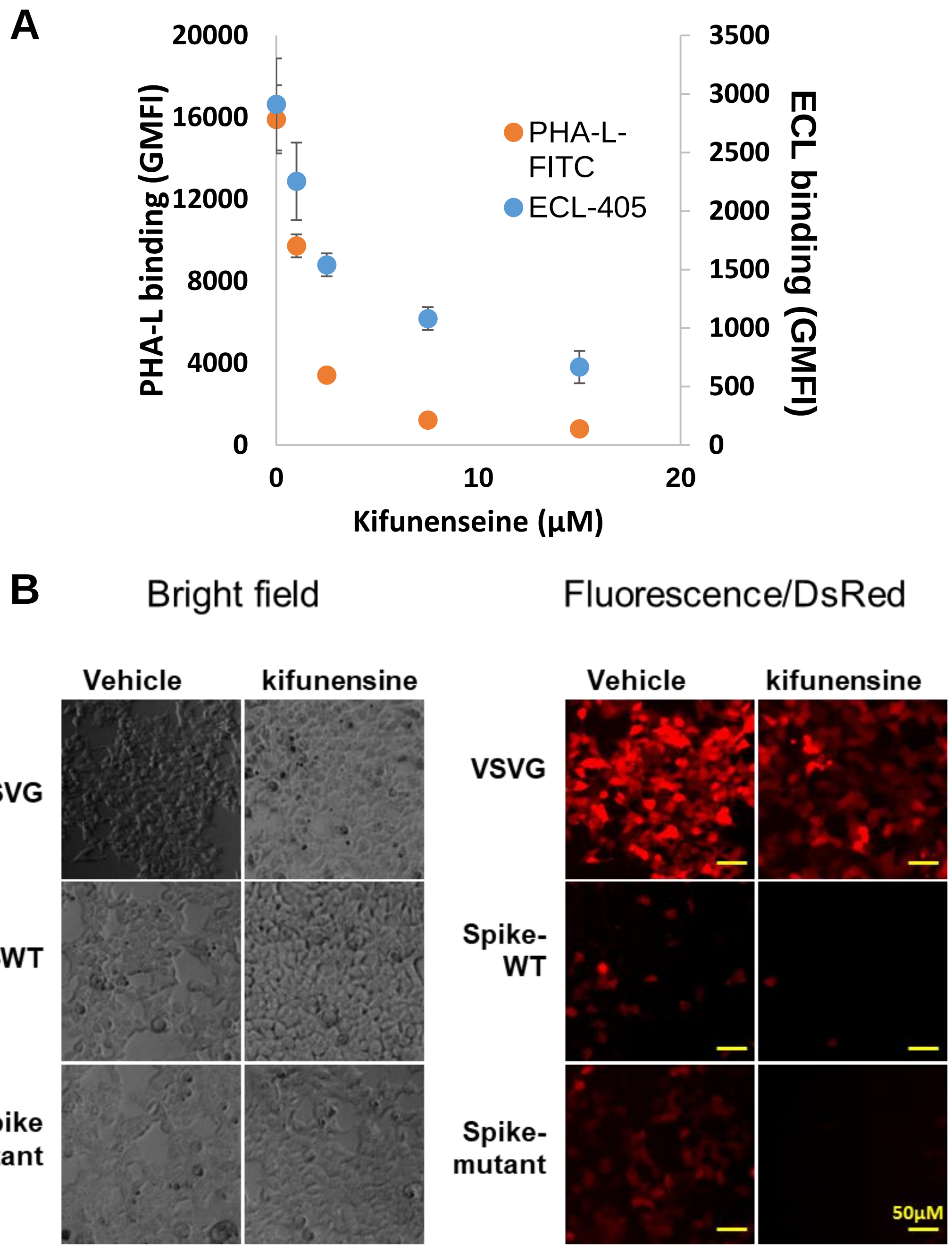


**Figure 5-figure supplement 1. Effect of kifunensine. A.** Lectin binding upon kifunensine treatment. HEK293T cells were cultured with varying concentrations of kifunensine or vehicle for 2 days. PHA-L binding measurement shows marked reduction in the cell surface expression of complex glycans and also a decrease in ECL engagement (recognizes lactosamine chains). Kifunensine did not affect cell growth rate or viability. It also did not reduce pseudovirus production titers. Data are Mean + STD (N=3). **B.** Kifunensine reduced Spike viral infection. Bright field images for cell monolayer (left) for microscopy data shown in main manuscript (reproduced in right panels). Kifunensine reduces Spike-WT and Spike-mutant pseudoviral entry by ~85-90%.
